# Enhanced post-traumatic headache-like behaviors and diminished contribution of peripheral CGRP in female rats following a mild closed head injury

**DOI:** 10.1101/865444

**Authors:** Dara Bree, Kimberly Mackenzie, Jennifer Stratton, Dan Levy

## Abstract

**Introduction:** Females are thought to have increased risk of developing posttraumatic headache (PTH) following a traumatic head injury, or concussion. However, the processes underlying this susceptibility remain unclear. We previously explored the development of PTH-like pain behaviors in a novel rat model of mild closed head injury, along with the ability of sumatriptan and an anti-calcitonin-gene-related peptide monoclonal antibody to ameliorate these behaviors. Here, we explored the development of PTH-like behaviors and the effectiveness of these headache therapies in females subjected to the same head trauma protocol.

**Methods:** Adult female Sprague Dawley rats were subjected to a mild closed head injury using a weight-drop device. Characterization of headache and pain related behaviors included assessment of changes in cutaneous cephalic and extracephalic tactile pain sensitivity, using von Frey monofilaments. Sensitivity to headache/migraine triggers was tested by examining the effect of systemic administration of a low-dose of glyceryl trinitrate (GTN). Treatments included acute systemic administration of sumatriptan and repeated systemic administration of a mouse anti-calcitonin-gene-related peptide monoclonal antibody. Serum levels of calcitonin-gene-related peptide were measured at various time points in females and males after the head injury.

**Results:** Female rats subjected to a mild closed head injury developed cutaneous mechanical hyperalgesia, that was limited to the cephalic region, and was resolved 4 weeks later. Cephalic pain hypersensitivity was ameliorated by treatment with sumatriptan, but was resistant to an early and prolonged treatment with the anti-CGRP monoclonal antibody. Following the resolution of the head injury-evoked cephalic hypersensitivity, administration of GTN produced a renewed and pronounced cephalic and extracephalic pain hypersensitivity that was inhibited by sumatriptan, but only partially by the anti-CGRP treatment. CGRP serum levels were elevated in females but not in males at 7 days post head injury.

**Conclusions:** Development of PTH-like pain behaviors following a mild closed head injury, and responsiveness to treatment in rats is sexually dimorphic. When compared to males, female rats display a prolonged state of cephalic hyperalgesia, increased responsiveness to a headache trigger, and a poorer effectiveness of an early and prolonged anti-CGRP treatment. The increased risk of females to develop PTH may be linked to enhanced responsiveness of peripheral and/or central pain pathways and a mechanism independent of peripheral CGRP signaling.

## Introduction

Post-traumatic headache (PTH) remains one of the most common and disabling symptoms following traumatic head injury. Defined as a secondary headache that develops within seven days of the head trauma (1), PTH often shares similar clinical characteristics with primary headaches, in particular migraine and tension-type headache (TTH) (2, 3). Currently, there exists no consensus on how PTH should be treated, primarily due to the poor understanding of its underlying mechanisms, which can be attributed in part to the paucity of well characterized and clinically relevant animal models with strong predictive validity. We recently characterized numerous PTH-like pain behaviors in male rats subjected to a mild closed head injury (mCHI) using a weight drop device (4). These include the development of acute cephalic allodynia, deficits in spontaneous exploratory activity and latent sensitization to the headache trigger glyceryl trinitrate (GTN). Furthermore, these pain behaviors were amenable to acute treatment with sumatriptan as well as repeated administration of an anti-calcitonin-gene-related peptide monoclonal antibody (Anti-CGRP mAb) starting immediately after the head injury (4), further strengthening the translational validity and potential utility of this animal model to study mechanisms of PTH.

Current understanding of PTH symptomatology and possible underlying mechanisms, as gleaned by clinical (3) and preclinical studies (5) are primarily based on findings in male subjects, in part due to the traditionally perceived idea of greater participation of males in activities with increased risk of head injuries and related concussions. However, recent clinical data suggest that females, in particular at adolescent age, have a similar or even higher risk for concussive injuries (6). Furthermore, females are now thought to have a similar or even increased risk, in particular at young age, of developing of PTH (7-10).

With mounting evidence of sex-differences in pain sensitivity and analgesic responses (11), as well as migraine-related mechanisms (12-14), the aim of the current study was to extend our knowledge of the pathophysiology of PTH by investigating potential sex-specific differences in the development or maintenance of PTH-like pain behaviors in females rats subjected to mCHI. In addition, we sought to study the relative efficacy of acute sumatriptan treatment as well as of an early and prolonged treatment with an anti-CGRP mAb.

## Materials & Methods

### Animals

All experiments were approved and conducted in compliance with the institutional Animal Care and Use Committee of the Beth Israel Deaconess Medical Centre, and the ARRIVE (Animal Research: Reporting of *in vivo* Experiments) guideline (15). Subjects were female rats (Sprague-Dawley rats, Taconic, USA, 8-9 weeks at the time of arrival). Animals were housed in pairs with food and water available *ad libitum* under a constant 12-hour light/dark cycle (lights on at 7:00 am) at room temperature. Studies were initialized after a week of acclimatization in the vivarium. All procedures and testing were conducted during the light phase of the cycle (08:00-15:00). Experimental animals were randomly assigned to either sham or mCHI as well as to the different pharmacological treatment groups.

### Experimental Mild Closed Head Injury (mCHI)

mCHI was induced using the weight-drop device as described previously in male rats (4). Briefly, rats were anesthetized with 3% isoflurane and placed chest down directly under a weight-drop concussive head trauma device. The device consisted of a hollow cylindrical tube (inner diameter 2.54 cm) placed vertically over the rat’s head. Mild closed head injury was induced by dropping a 250 g projectile through the tube from a height of 80 cm, striking the center of the head. To ensure consistency of the hit location, animals were placed under the weight drop apparatus so that the weight struck the scalp slightly anterior to the center point between the ears. A foam sponge (thickness 3.81 cm, density 1.1 g/cm^3^) was placed under the animals to support the head while allowing some linear anterior-posterior motion without any angular rotational movement at the moment of impact. A repeated strike was prevented by capturing the weight after the first strike. All animals regained their righting reflex within 2 min (which likely reflects the recovery from anesthesia). Immediately after the impact, animals were returned to their home cages for recovery and were neurologically assessed up to 7 days post-injury for any behavioral abnormalities suggestive of a major neurological (i.e. motor) impairment. Sham animals were anesthetized for the same period of time as mCHI animals, but not subjected to the weight drop. There was 0% mortality after the mCHI procedure with no evidence of skull fractures or cortical bleeding.

### Open field behavior

Activity Monitor (Med Associates, Vermont, USA) was used to assess mCHI-evoked changes in locomotor activity, exploratory behavior, and anxiety-like behavior in an open field arena (43 × 43 × 30 cm), as previously described (4). The monitoring system records the movement of animals in the horizontal (X-Y axis) and vertical (Z-axis) planes using 16 infrared beams and detectors, spaced 2.54 cm apart. Data analyzed included total distance moved, vertical rearing events (exploratory behavior), and relative time spent in the center of the arena (30 cm × 30 cm) during a 20 minutes session. The arena was lit with a single white LED bulb on a dimmer switch to maintain a homogenous lighting across the arenas (80 lux). The arena was cleaned with a mild detergent and dried to remove odor cues between successive rats.

### Assessment of tactile pain hypersensitivity following mCHI

Behavioral tests were performed during the light phase (08:00-15:00). The method was previously used by us and others to study PTH- and migraine-related pain behaviors (4, 16-18). Briefly, animals were placed in a transparent flat-bottomed acrylic holding apparatus (20.4 cm × 8.5 cm). The apparatus was large enough to enable to the animals to escape the stimulus. Animals were habituated to the arena for 15 minutes prior to the initial testing. In order to determine if the animals developed pericranial (cephalic) tactile hypersensitivity, the skin region, including the midline area above the eyes and 2 cm posterior, was stimulated with different von Frey (VF) filaments (0.6 g– 15 g/force) (18011 Semmes-Weinstein Anesthesiometer kit). Development of hind paw hypersensitivity was tested by stimulating, using the VF filaments, the mid-dorsal part of the hind paw. We evaluated changes in withdrawal thresholds, as well as non-reflexive pain responses to the stimulation using a method previously described in this and other headache models (19-21) by recording 4 behavioral responses adapted from Vos et al. (22) as follows: 0) *No response*: rat did not display any response to stimulation 1) *Detection*: rat turned its head towards stimulating object and latter is explored usually by sniffing; 2) *Withdrawal*: rat turned its head away or pulled it briskly away from stimulating object (usually followed by scratching or grooming of stimulated region); 3) *Escape/Attack*: rat turned its body briskly in the holding apparatus in order to escape stimulation or attacked (biting and grabbing movements) the stimulating object. Starting with the lowest force, each filament was applied 3 times with an intra-application interval of 5 seconds and the behavior that was observed at least twice was recorded. For statistical analysis, the score recorded was based on the most aversive behavior noted. The force that elicited two consecutive withdrawal responses was considered as threshold. To evaluate pain behavior in addition to changes in threshold, for each rat, at each time point, a cumulative pain score was determined by combining the individual scores (0–3) for each one of the VF filaments tested. All tests were conducted and evaluated in a blinded manner.

### mCHI evoked latent sensitization to GTN

The development of mechanical hyperalgesia in response to systemic administration of a previously subthreshold dose of the headache trigger glyceryl trinitrate (GTN, 100μg/kg i.p., American Reagents, USA) when animals returned to baseline following mCHI was assessed as described (4, 21). After obtaining pre-GTN baseline cephalic and hind paw VF responses at day 29 post mCHI/sham, animals received GTN on day 30 and were assessed for changes in cephalic and hind paw mechanical pain thresholds 4 hours later.

### Pharmacological treatments

Sumatriptan (Tocris, USA) was freshly dissolved in 0.9% saline and administered intra-peritoneal (i.p.) at a dose of 1mg/kg in a volume of 1ml/kg. Drug dose and times of administration were based on the pharmacokinetics of the drugs, studies demonstrating their efficacy in animal and human models of trigeminal pain (16, 23), and our previous data on PTH-like behaviors following mCHI in males (4). Anti-CGRP mAb and its corresponding isotype IgG were provided by TEVA Pharmaceuticals, and formulated in phosphate buffered saline (PBS). mAb and IgG were injected i.p at a dose of 30 mg/kg, immediately after the head injury and every 6 days subsequently up to day 30. This dosing regimen has been shown previously to alleviate PTH-like pain behavior (i.e. cephalic allodynia) following mCHI in male rats (4) and pain behaviors in other chronic migraine models (24).

### Peripheral blood collection and serum CGRP analysis

At multiple time points (baseline and 72h, Day 7, Day 14 post mCHI) rats were lightly anesthetized and placed chest down on a heating blanket. The tail was dipped in warm water for 10 seconds in order to increase vascular flow and swabbed with alcohol. Approximately 300 microliters of blood was collected from the tail vein using a BD Vacutainer Safety-Lok blood collection needle (Becton Dickinson, USA) and placed in a microfuge tube containing 10 microliters of protease inhibitor (Millipore Sigma). Immediately after collection, the blood was placed on ice to coagulate for 30 minutes. Following coagulation, blood was centrifuged at 3000g for 10 min at 4 degrees. The serum supernatant was transferred to fresh microfuge tubes, snap frozen on dry ice and stored at – 80 degrees Celsius. Serum samples were analysed for CGRP levels via a custom ELISA. Briefly, a 96-well plate was coated with a mouse anti-CGRP capture antibody (Bertin Bioreagent) and incubated overnight at 4°C. The coated plate was washed three times with wash buffer (PBS with 0.05% Tween 20) and then blocked using the Sword Blocker SBL-501 reagent (Sword Diagnostics) for 1 hour at room temperature. After several washes, serum samples and a range of CGRP standards diluted in Sword Diluent SDI-802 were pipetted into the coated plate with a human anti-CGRP detection antibody (Teva Pharmaceuticals) and incubated at room temperature for 2 hours. After washing the plate, captured CGRP analytes were complexed to a HRP-conjugated mouse anti-human antibody (Southern Biotech) at room temperature for 1 hour. Following several washes, Sword detector reagents consisting of a substrate/peroxidase mixture were used according to the manufacturer’s instructions. Resonance Raman signals generated using Sword reagents were measured using fluorescence intensity detection at an excitation/emission wavelength of 530nm/730nm using a BioTek Cytation 5 Microplate Reader.

### Data analyses

Statistical analyses were conducted using Graph pad prism (version 8.0). All data are presented as the means ±standard error of the mean. Mechanical pain threshold data, obtained using VF filaments was log-transformed, which is more consistent with a normal distribution (25). To study time-course changes in open-field behaviour, and responses to mechanical stimuli, data was analysed using a mixed design ANOVA to determine the effects of time and treatment. Data included passed the Brown-Forsythe test, indicating equal variance. We used Fisher’s LSD post hoc tests, and correction for multiple comparisons was conducted using the Benjamini and Hochberg false discovery rate (FDR) controlling procedures (26). Responses to GTN were assessed using two-tailed paired t-test. The effects of drug treatment on the changes in nociceptive behaviours were analysed using two-tailed unpaired t-test. Changes in CGRP plasma levels were tested using one-Way Brown-Forsythe ANOVA test. Significant p and q value were set as 0.05.

## Results

### mCHI-evoked changes in open-field behavior

Previously, we observed that mCHI did not affect the overall distance moved of male rats in an open field, indicating no gross motor deficits, but led to a reduction in rearing activity suggesting deficits in exploratory behavior, potentially due to a mild TBI (4, 27, 28). Here, we observed that females subjected to a similar mCHI protocol also did not exhibit changes in distance moved when tested up to 14 days post mCHI (Time, F_4,56_ = 2.1, p = 0.1; Treatment, F_1,14_ = 0.4, p=0.5; **Figure 1(b)**). Similarly, mCHI females also did not exhibit any change in rearing activity (Time, F_4,56_ = 2.4; p = 0.07; Treatment F_1,14_ = 0.1, p=0.7; **Figure 1(c)**) pointing to the possibility of a milder form of TBI when compared to males. The lack of a noticeable decrease in exploratory rearing activity, however, may be due to a lower baseline rearing behavior in our female cohort when compared to the matching male cohort (females; 109.5±8.4 beam breaks/20 min vs males; 167.9 ±12.1 beam breaks/20 min; p < 0.001, unpaired t-test; not shown). Females subjected to mCHI, however, displayed reductions in center zone exploration, indicating increased thigmotaxis (Time, F_4,56_ = 7.9; p < 0.0001; Treatment, F_1,14_ = 4.63; p < 0.05; **Figure 1(d)**). Post-hoc analysis revealed decreased center zone exploration at 2 and 3 days following mCHI (q<0.01 for both) suggesting an acute increase in anxiety-related behaviors, potentially related to mTBI (29, 30). The increased thigmotaxis response in females was likely a specific sex-dependent response to the mCHI given that males and females exhibited similar pre-mCHI baseline values (29.1±4.2% of time spent in center vs 32.3 ± 2.5 % of time spent in center; p = 0.52, unpaired t-test, not shown).

**Figure 1:**
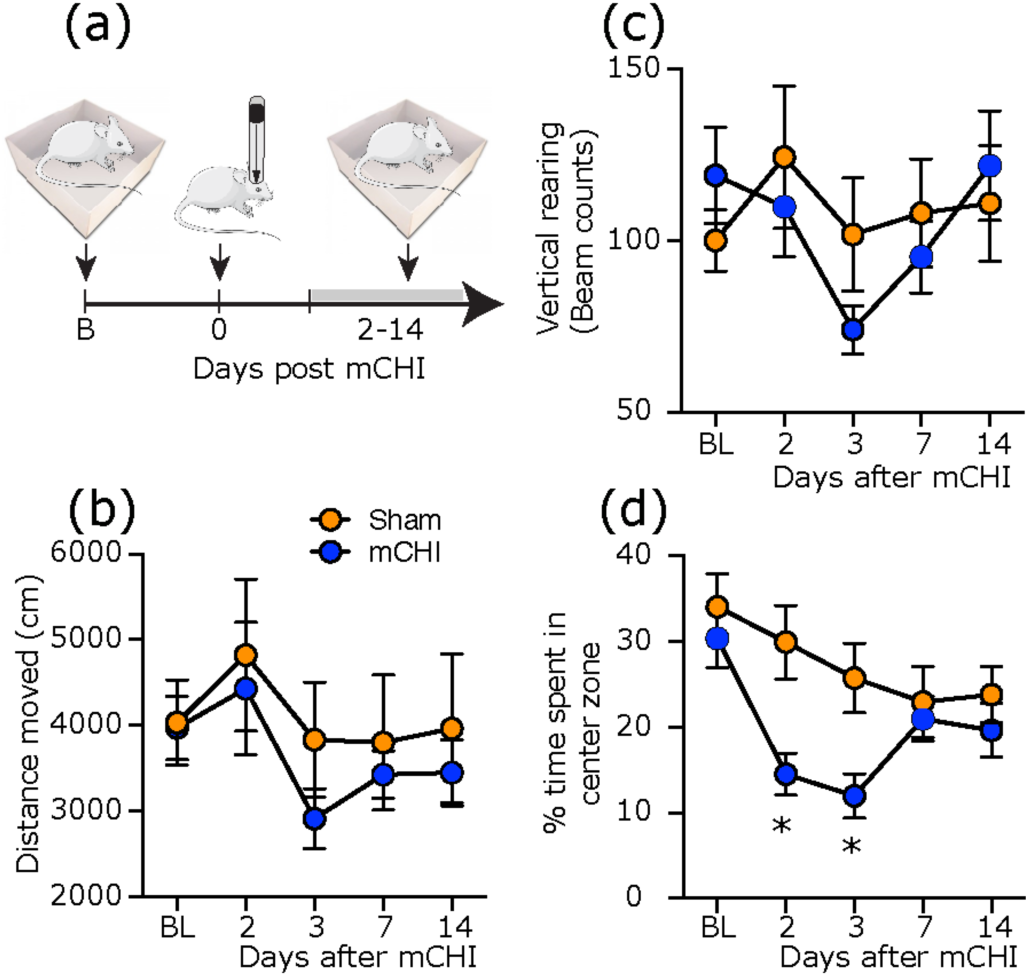
Changes in open field behaviors in females following mCHI. (a) Scheme of the experimental design. Rats were subjected to a baseline open-field testing, following by mCHI, and additional testing 2-14 days later. (b) Changes in distance moved, (c) exploratory vertical rearing, and (d) % time spent zone. Two-way repeated measures ANOVA, followed by post-hoc test between mCHI and sham animals indicate an acute decrease in time spent in center zone (and increased thigmotaxis) at 2 and 3 days following mCHI with no changes in other open-field parameters. Data are mean and SEM (n=8). * q< 0.05 (FDR-corrected values after Fisher’s least significant difference post-hoc test at selected time points vs similar time points in sham control animals). FDR, false discovery rate; mCHI, mild closed-head injury.

### Development of mechanical pain hypersensitivity following mCHI

Male rats subjected to mCHI develop cephalic mechanical hypersensitivity that resolves by day 14 post mCHI. Here, we observed that females subjected to a similar mCHI protocol, also exhibited decreased cephalic mechanical thresholds when compared to sham animals (Time, F_6,129_ = 15.3, p < 0.001; Treatment: F_1, 27 =_ 25.8, p < 0.001; **Figure 2(b)**). The duration of this allodynic response was, however, longer than we previously observed in males, and was resolved only by day 29 post mCHI. Females subjected to mCHI also exhibited an increase in pain score in response to cephalic mechanical stimulation (Time: F_6,131_ = 10.3, p<0.0001; Treatment: F_1, 27 =_ 9.3, p < 0.001; **Figure 2(C)**); this hyperalgesic behavior, however was shorter lasting and resolved by day 14 post mCHI. As in males, the development of cephalic pain hypersensitivity was not accompanied by extracephalic changes throughout the 29 days observation period (threshold, F_6,137_ = 8.8, p<0.001 for time; F_1,27_ = 1.4, p = 0.25 for treatment; pain score, F_6,137_ = 9.5, p<0.001 for time; F_1,27_ =1.4, p = 0.24 for treatment, **Figures 2(d and e)**).

**Figure 2:**
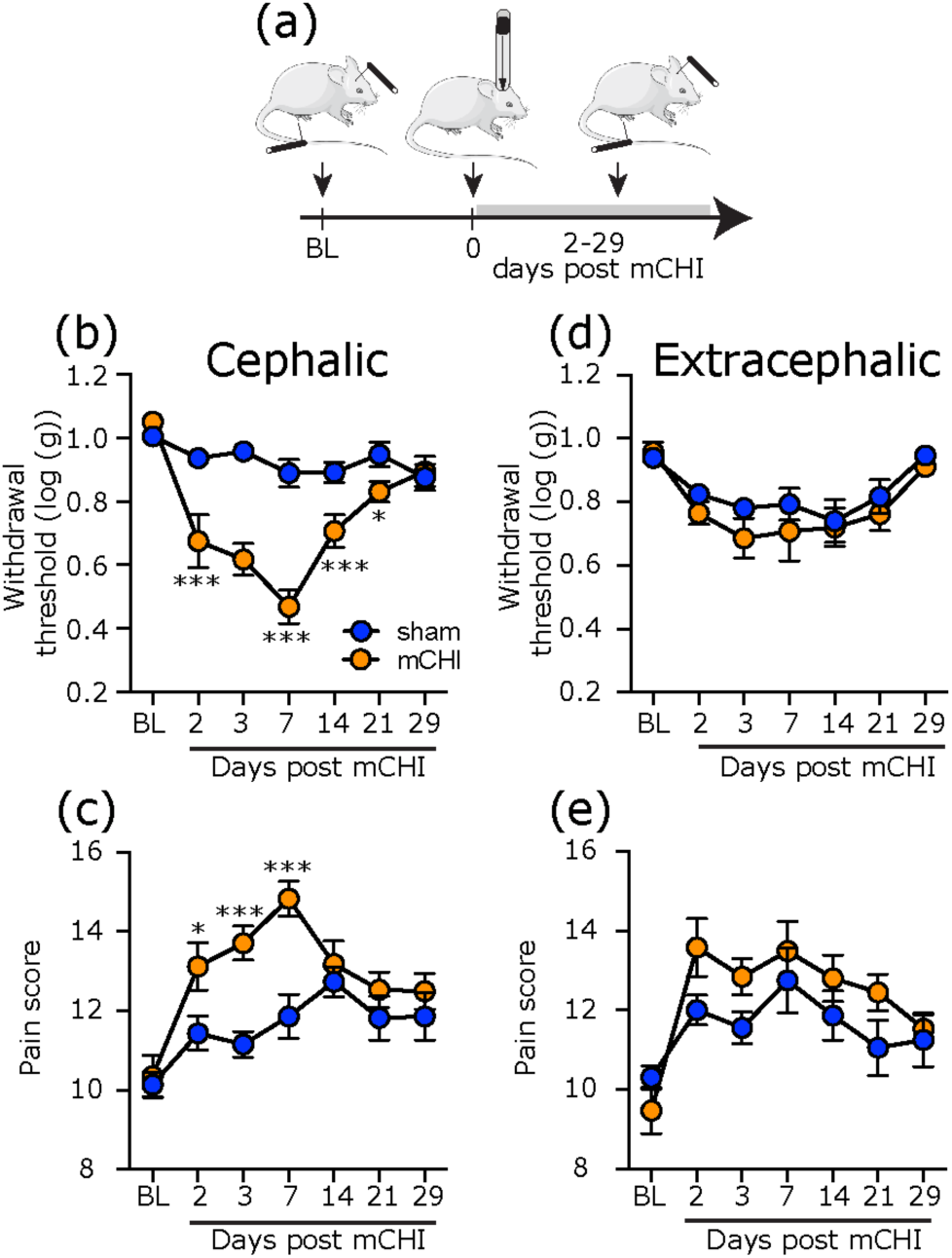
Development of prolonged cephalic cutaneous mechanical hypersensitivity in females following mCHI. (a) Schematic of experimental design. Rats underwent baseline von Frey testing of mechanical pain sensitivity at the cephalic and extracephalic (hind paw) regions, followed by mCHI, and further nociceptive testing at these locations 2-29 days later. Time course changes in cephalic (b) and extracephalic (d) mechanical pain withdrawal thresholds and corresponding cumulative pain scores at the cephalic (c) and extracephalic (e) regions. Two-way repeated measures ANOVA, followed by post-hoc test between mCHI and sham animals indicate a prolonged decrease in withdrawal thresholds and increase in pain score following mCHI at the cephalic region, but no extracephalic changes. Sham: n=12; mCHI: n=17. * q<0.05, *** q<0.001 (FDR-corrected values after Fisher’s least significant difference post-hoc test at selected time points vs similar time points in sham control animals).

### mCHI-evoked prolonged latent cephalic and extracephalic sensitization to GTN

Male rats subjected to mCHI exhibit latent sensitization manifested as a cephalic hyperalgesic response to a subthreshold dose of the headache trigger GTN, after the recovery from the acute hyperalgesic phase (4). Here, we observed latent sensitization in females that was similar with regard to the cephalic response, but also involved the extracephalic region. When compared to baseline values obtained on day 29 post mCHI, administration of GTN on day 30 resulted in cephalic mechanical hypersensitivity (p<0.001, for thresholds and pain scores, **Figures 3(b and c)**), an effect that was not observed in sham animals. Similarly, GTN-evoked hind paw hypersensitivity in mCHI females (p < 0.001, for both threshold and pain score (**Figures 3(d, and e)**), but not in females subjected to a sham procedure.

**Figure 3:**
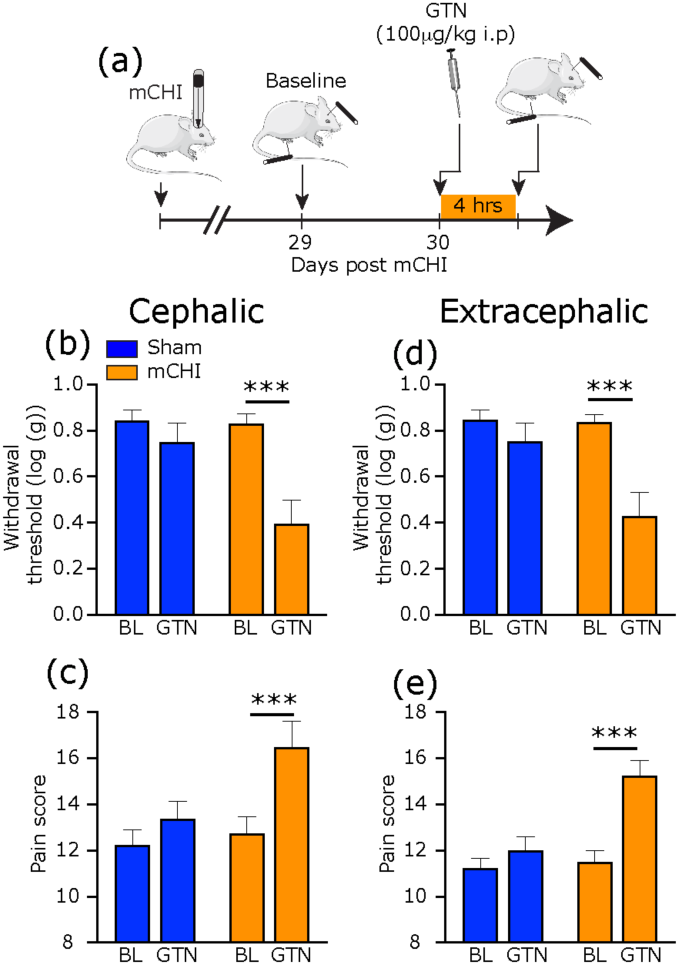
Cephalic and extracephalic sensitization to GTN following in females 30 days following mCHI. (a) Scheme of the latent sensitization experimental design. mCHI rats were subjected to baseline behavioral testing at the cephalic and extracephalic regions on day 29 after mCHI (pre-GTN baseline), and then a day later (day 30), 4 hours after systemic administration of subthreshold dose of GTN. Summary of the cephalic withdrawal thresholds (b) and pain scores (c) in sham and mCHI animas at baseline and following GTN treatment. GTN-evoked extracephalic changes are illustrated in (d) and (e). GTN evoked significant mechanical pain hypersensitivity at both regions tested. Bar data are means±SEM, Sham; n=8; mCHI: n=8; *** p<0.001, two-tailed paired t-test.

### Acute sumatriptan treatment ameliorates mCHI-induced cephalic hypersensitivity, and GTN-evoked cephalic and extracephalic hypersensitivity

In males, acute treatment with sumatriptan alleviates the cephalic mechanical hypersensitivity following mCHI, and the cephalic pain hypersensitivity in response to GTN (4). Here, we found that acute sumatriptan treatment in females, at 7 days post-mCHI, exerted a similar anti-hyperalgesic effect by decreasing the cephalic pain threshold (p<0.001, **Figure 4(b)**), as well as the associated pain response (p <0.01, **Figure 4(c)**). Acute sumatriptan treatment was also effective at inhibiting the GTN-evoked decrease in cephalic mechanical pain threshold and increased pain score (p<0.001, p<0.05 respectively, **Figures 5(b and c)**). Sumatriptan was also able to inhibit the GTN-evoked decrease in hind paw mechanical pain threshold, and increased pain score (p<0.001, p<0.001, **Figure 5 (d and e)**).

**Figure 4:**
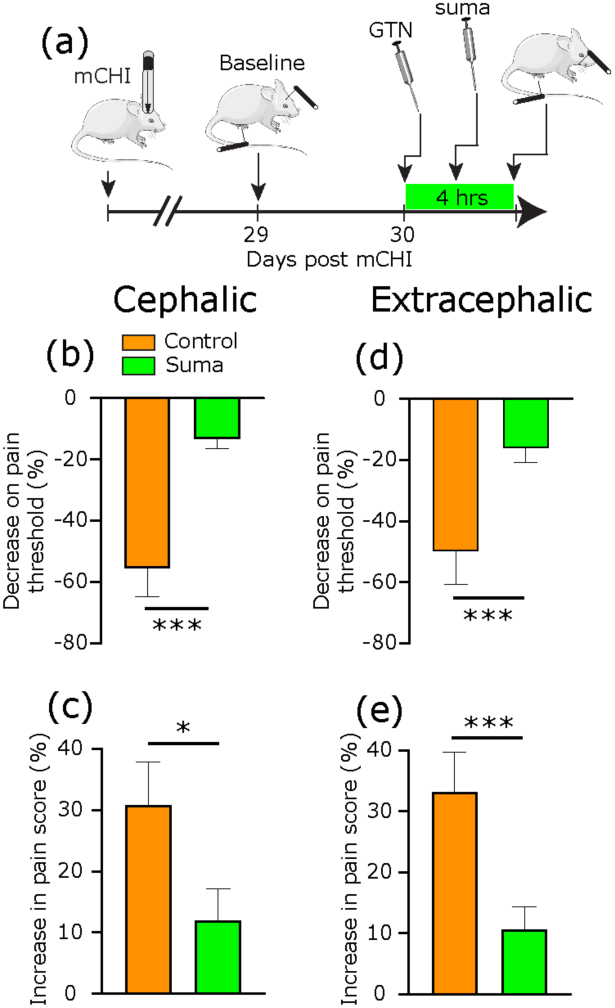
Acute sumatriptan treatment exert an anti-hyperalgesic effect in mCHI females. (a) Schematic of experimental design. Rats underwent baseline von Frey testing of cephalic mechanical pain sensitivity, followed by mCHI. On day 7 post mCHI, animals were treated with sumatriptan or vehicle, and then subjected to nociceptive testing 2 hours later. Summary of sumatriptan-related changes in withdrawal thresholds (b) and related pain scores (c). Bar data are means±SEM, Control; n=10; Suma n=10, *** p<0.001, unpaired t-test.

**Figure 5:**
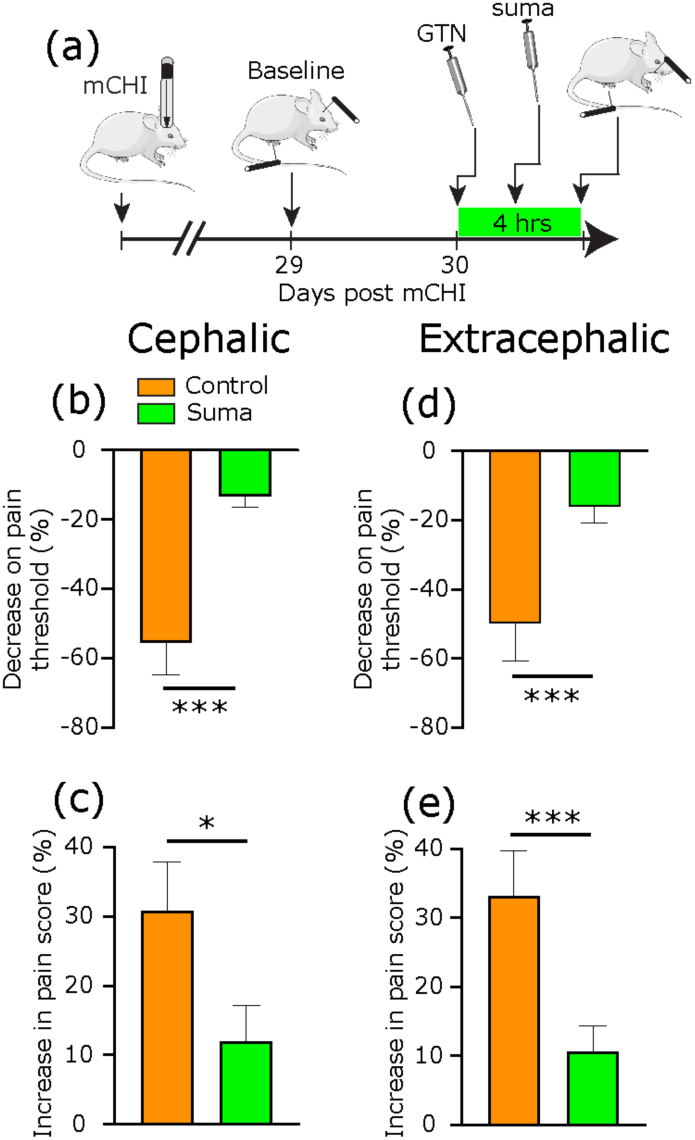
Acute sumatriptan treatment exerts an anti-hyperalgesic effect on GTN-evoked cephalic and extracephalic mechanical pain hypersensitivity in mCHI females. (a) Schematic of experimental design. mCHI rats were subjected to baseline behavioral testing at the cephalic and extracephalic regions on day 29 after mCHI (pre-GTN baseline), and then a day later (day 30). Sumatriptan was administered 1 hour prior to systemic administration of subthreshold dose of GTN followed by von Frey testing 4 hours later. Summary of GTN-evoked changes in cephalic withdrawal thresholds (b) and related pain scores (c). Summary of GTN-evoked changes at the extracephalic site. Means±SEM, Control: n=8; Suma: n=9; *** p<0.001; * p<0.05; unpaired t-test.

### Early and repeated administration of anti-CGRP mAb does not prevent mCHI-induced cephalic mechanical hypersensitivity, but partially inhibits GTN-evoked pain hypersensitivity in female rats

In males, repeated administration of anti-CGRP mAb, starting immediately following the mCHI, inhibits the cephalic hypersensitivity and the prolonged latent cephalic sensitization to GTN (4). Here, in contrast, when compared to treatment with a control IgG, similar administration of the anti-CGRP mAb in females did not block the decrease in cephalic pain threshold (Time, F_3,90_ = 17.1, p < 0.0001; Treatment: F_1, 30_ = 0.1, p = 0.7, **Figure 6(b)**), or the associated increase in pain score (Time, F_3,90_ = 41.5, p < 0.0001; Treatment: F_1, 30 =_ 0.7, p = 0.42, **Figure. 6(c)**). Anti-CGRP treatment was also less effective in inhibiting the hyperalgesic response to GTN at 30 days following mCHI. When compared to treatment with the control IgG, anti-CGRP mAb inhibited the GTN-evoked decrease in cephalic thresholds (p<0.05, **Figure 7(b)**), but not the associated increase in pain score (p = 0.23, **Figure 7(c)**). Treatment with the anti-CGRP mAb was also ineffective in inhibiting the GTN-evoked hind paw hypersensitivity, when compared to IgG (p = 0.1 for threshold changes; p = 0.36 for pain score changes, **Figures 7(d and e)**).

**Figure 6:**
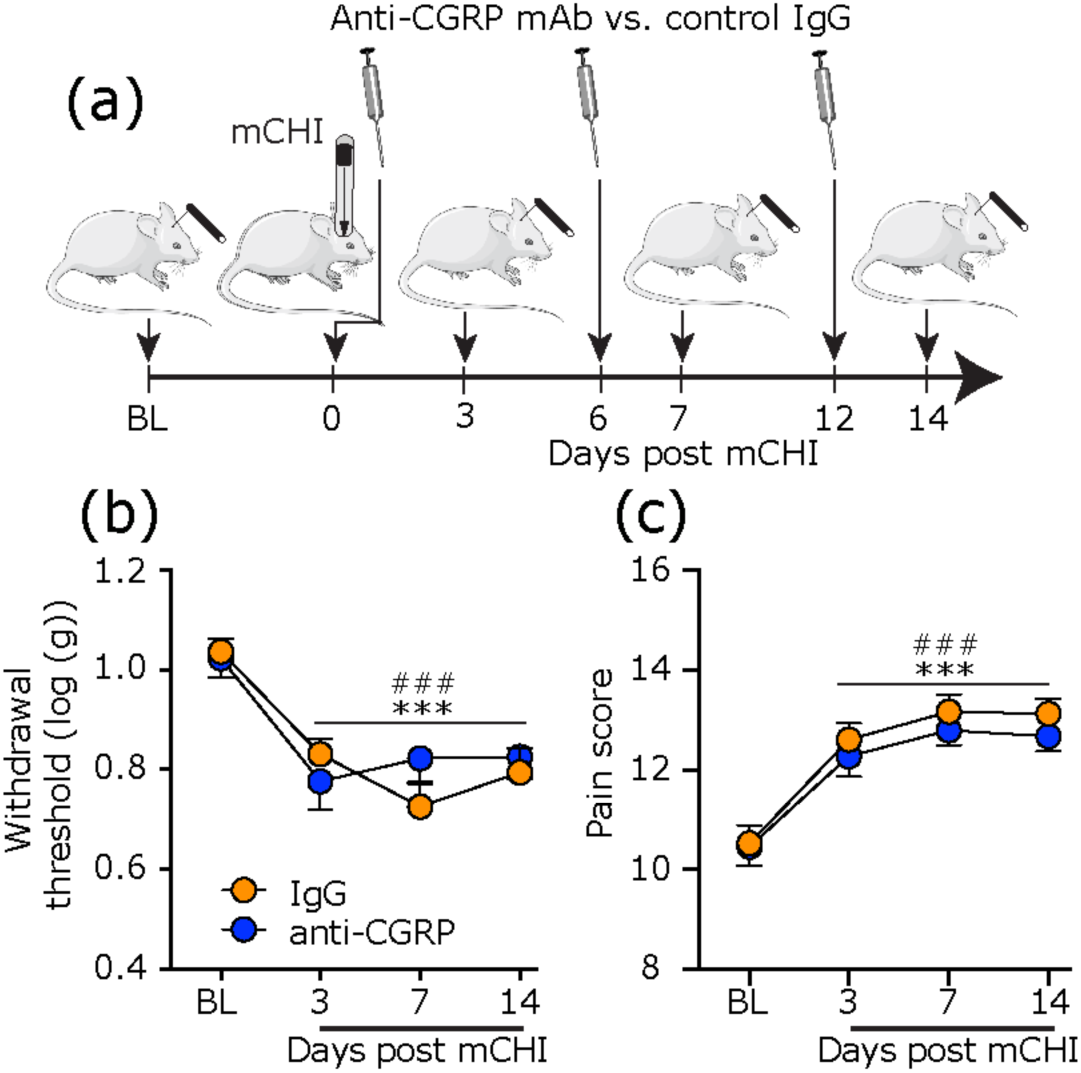
Treatment with anti-CGRP mAb does not affect the development of cephalic pain hypersensitivity in mCHI females. Schematic of experimental design. Rats underwent baseline von Frey testing of cephalic mechanical pain sensitivity, followed by mCHI. Anti-CGRP mAb or a control IgG were administered immediately after the mCHI and then every 6 days. Two-way repeated measures ANOVA between anti-CGRP and IgG treatments revealed no effect of treatment on the decrease in cephalic withdrawal threshold (b), or the increase in pain score (c). Means±SEM, Anti-CGRP: n=16; control IgG: n=16; *** q<0.001; ### q< 0.001 (FDR-corrected values after Fisher’s least significant difference post-hoc test at 3-14 days post mCHI vs baseline for anti-CGRP and IgG respectively).

**Figure 7:**
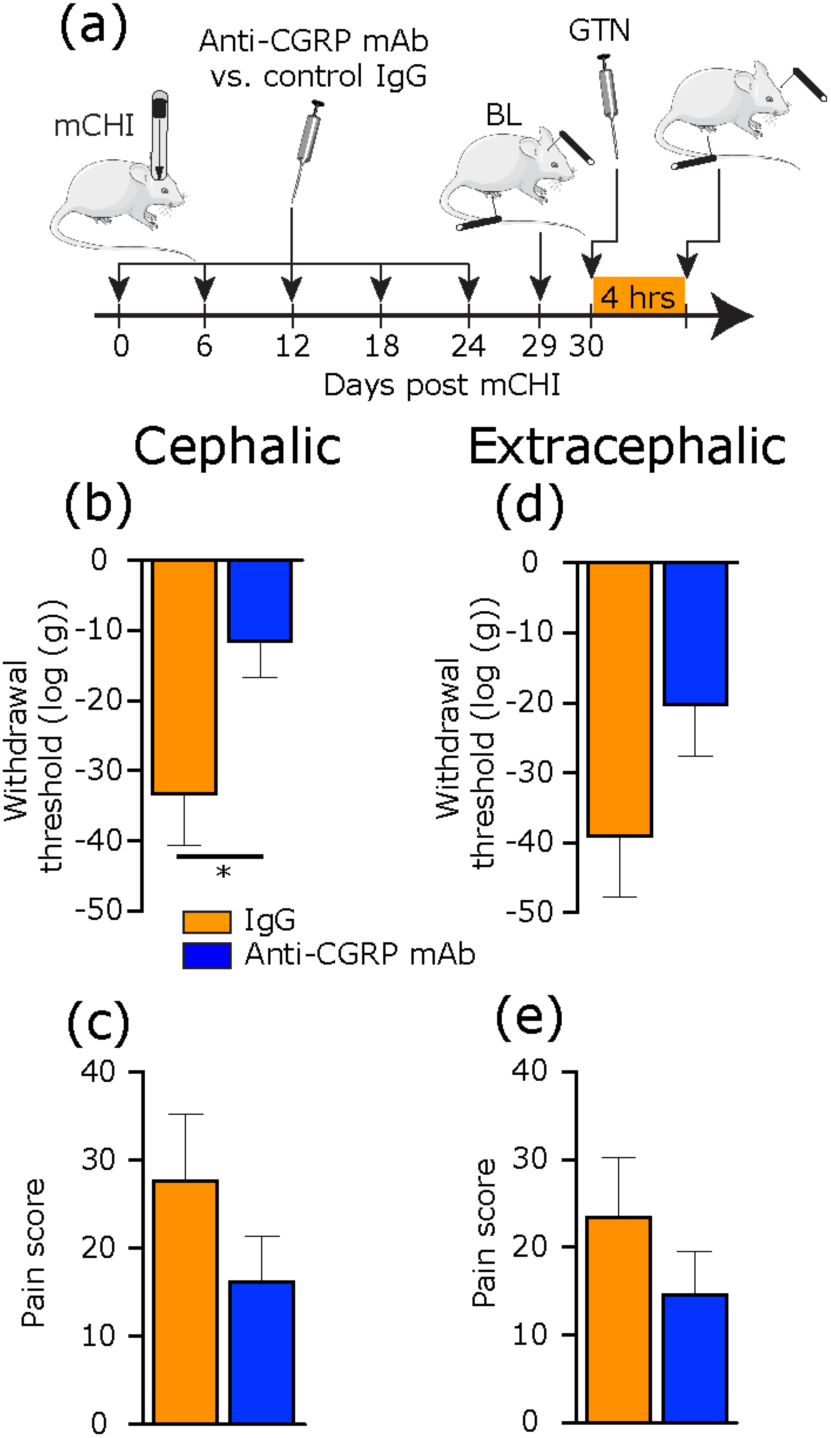
Treatment with anti-CGRP mAb partially inhibits latent sensitization to GTN in mCHI females. (a) Schematic of experimental design. mCHI rats received treatment with anti-CGRP mAb or a control IgG immediately after the mCHI and then every 6 days. GTN-evoked cephalic and extracephalic mechanical pain hypersensitivity was assessed at days 29 (baseline) and day 30 (GTN administration). Summary of the GTN-evoked changes in cephalic withdrawal thresholds (b) and pain scores (c). GTN-evoked extracephalic changes are illustrated in (d) and (e). Means±SEM, Anti-CGRP: n=12; IgG: n=12; * p<0.05 unpaired t-test.

### mCHI in female leads to transient increase in serum CGRP levels

Having found a lack of inhibitory effect of the anti-CGRP mAb on the mCHI-evoked pain hypersensitivity and a partial effect on the latent sensitization to GTN in females, we examined serum CGRP levels in females at baseline and up to 14 days following mCHI. Overall, we found time-related increases in the CGRP serum levels following mCHI (W_3,18.9_ = 4.5, p<0.01). Post-hoc analyses revealed an increase in CGRP serum levels only at day 7 (q<0.001, **Figure 8(b)**). We also examined serum CGRP levels in male rats, in which treatment with anti-CGRP mAb produced an anti-hyperalgesic effect following mCHI (4). However, we did not detect any time-course changes in serum CGRP levels for up to 14 days following mCHI (W_3,15.9_ = 0.83, p = 0.49).

**Figure 8:**
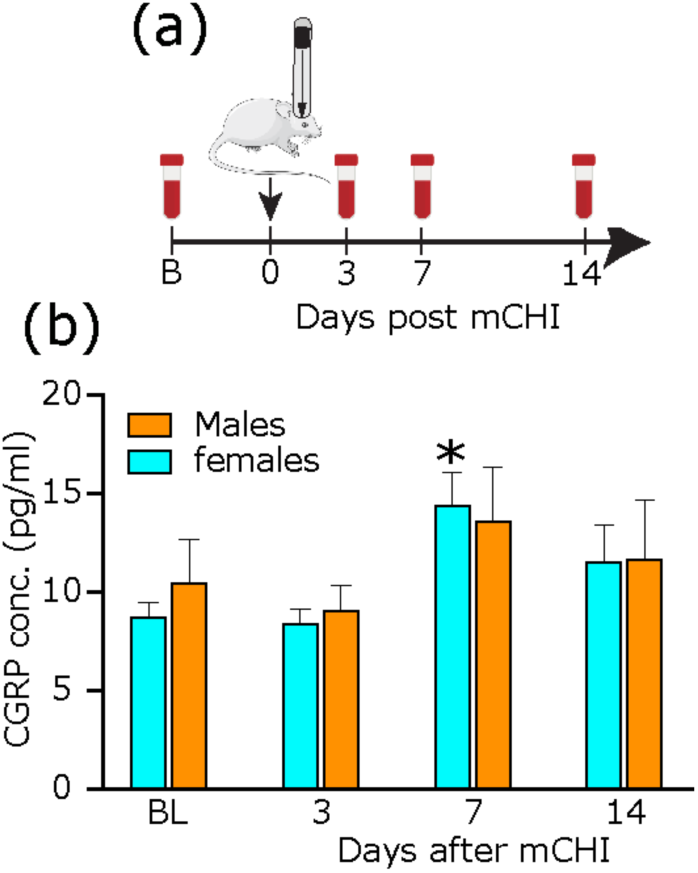
Sex-related changes in serum CGRP levels following mCHI. (a) Schematic of experimental design. Serum CGRP levels were assessed in blood samples collected at baseline, and 3,7, and 14 days post mCHI. (b) Summary of the Serum CGRP levels in females and males. Data are Means±SEM, number of animals at each time point are in parentheses; * q< 0.05 (FDR-corrected values after Fisher’s least significant difference post-hoc test of mCHI vs baseline).

## Discussion

Building upon our previous work in male rats subjected to mCHI (4), the present study aimed to characterize changes in open-field behavior, and PTH-like pain behaviors following a similar traumatic head injury protocol in female rats. Our data point to key sex differences, including, enhanced pain responses, an acute increase in anxiety-like behavior, and decreased responsiveness to anti-CGRP mAb treatment in females.

A key sensory change identified in male subjects suffering from PTH (31), and in male rats subjected to mCHI (4) is the presence of cutaneous cephalic mechanical pain hypersensitivity. The development of cephalic mechanical allodynia is thought to reflect the sensitization of trigeminal pain pathways, a process that contributes to the pain in other types of headache, in particular migraine (32). Our current data suggests that mCHI in females leads to an enhanced cephalic hyperalgesic response, in particular a much longer duration, when compared to males subjected to a similar head injury protocol. Female sex has been suggested as a key positive predictor for development of PTH (33-35), and has been attributed to their higher prevalence of pre-existing headache and migraine (33, 34). Given that our rats did not encounter any potential headache producing event prior to the mCHI, our findings rather point to other/additional mechanisms, potentially related to increased nociceptive processing at the level of the peripheral and/or the central nervous system in females (14, 36-38), in particular facilitation of the process underlying the development of trigeminal central sensitization in response to head trauma.

Another key finding of the current study was the inability of an early and sustained anti-CGRP mAb treatment to ameliorate the prolonged cephalic mechanical hypersensitivity post-mCHI in females, in contrast to what we previously observed in males (4). Of importance, was the lack of the anti-hyperalgesic effect of the anti-CGRP mAb was despite the elevation of serum CGRP levels post-mCHI, and strongly suggests that the mCHI-evoked cephalic hypersensitivity in females may not involve peripheral CGRP signaling. It is noteworthy that we did not detect a similar increase in serum CGRP levels in males subjected to mCHI, which is in agreement with a previous study that employed a similar mCHI approach (39). The inconsistency between the effectiveness of anti-CGRP mAb treatment in this model and the changes in serum CGRP levels suggests that the latter may be a poor indicator of peripheral CGRP involvement in mediating a headache-like behavior. Our current finding that acute sumatriptan treatment was effective in reducing the cephalic hyperalgesic behavior in mCHI females, as previously reported in males (4), suggests that its therapeutic mode of action in this model may not involve alteration of the peripheral CGRP signaling cascade, and is likely to bypass the processes that is influenced by sex differences, as observed in other rodent models of persistent migraine-like cephalic pain hypersensitivity (36, 40, 41).

In addition to the prolonged cephalic hyperalgesic response in mCHI females, the enhanced pain response in females was also manifested by the development of latent sensitization to GTN that involved the cephalic region, as in males, but also an extracephalic hyperalgesic response to GTN, unlike in males. This increased responsiveness of females to GTN is reminiscent of the greater delayed extracephalic hyperalgesia observed in females following systemic administration of GTN at a higher dose, or after peripheral administration of a low dose (41, 42), and may be linked to a migraine-related mechanism that is influenced by sex differences. In males subjected to mCHI, the development of latent sensitization to GTN involves an immune-related mechanism, linked to mast cells (21). The previous findings that mCHI involves the degranulation of meningeal mast cells (43) and that meningeal mast cell density in females is higher than in males (13), points to the possibility that meningeal mast cells may play a role in the enhanced nociceptive response in females following mCHI, and particularly the prolonged latent sensitization.

Our finding in females that early and prolonged treatment with the anti-CGRP mAb was only partially effective in ameliorating the latent cephalic sensitization, unlike in males, suggests that this process may not be fully dependent on peripheral CGRP signaling, and that it is also under the influence of a sex-dependent mechanism. It is noteworthy that the lack or incomplete anti-hyperalgesic effect of anti-CGRP mAb treatment in females in this PTH model is in contrast to previous studies of migraine-related behaviors linked to peripheral CGRP signaling, which reported either lack of sex differences (44, 45), or female specific pro-nociceptive effects (14). Taken together, our data points to a potential differential role for peripheral CGRP signaling in the pathophysiology of PTH and migraine at least in females. The possibility that enhanced and persistent secretion of CGRP from injured meningeal afferent endings that innervate the subarachnoid space (i.e. leptomeningeal afferents), and which could not be targeted by peripherally administered mAb, plays a key role in PTH in females should also be considered.

The finding that sumatriptan treatment was effective in blocking the GTN-evoked extracephalic hypersensitivity, while anti-CGRP mAb was completely ineffective, points to the possibility that a central, rather than peripheral nociceptive process is involved in mediating PTH females. However, sumatriptan may also act peripherally, potentially via a vascular mechanism, to inhibit the GTN-evoked hyperalgesic response (42). Head trauma and associated brain injury give rise to peripheral changes, including altered immune response (46), and additional studies will be required to determine the relative contribution of mCHI-evoked peripheral vs central changes that mediate the enhanced extracephalic response to GTN.

A potential contributing factor to the enhanced response of females in this head trauma model is their relative smaller size when compared to males at the time of injury. At present we cannot exclude the possibility that the extended duration of the hyperalgesic response and enhanced hyperalgesic responsiveness to GTN we observed in females involved a greater intracranial damage with extended brain injury. However, the lack of changes in open field activity, including exploratory behavior, which agrees with other TBI-related behaviors noted in similar models of head trauma (47) argues against such a possibility. The development of acute anxiety-like behavior (i.e. thigmotaxis) in females, which was not detected previously in mCHI males, could possibly point to an enhanced secondary condition of brain injury (47). However, since increased thigmotaxis is also observed in other pain models (48-50), it may also signify the interdependent relationship between pain and anxiety. The present study did not monitor the estrous cycle or related hormonal changes, which may have influenced the data. However, the variability noted in the post mCHI data in females was reminiscent of that observed in our previous male study, and is in agreement with finding from a large data set of physiological, and behavioral measures (51), suggesting that estrous cycle-related changes in the levels of circulating hormones in females may not play a large role in this model.

## Conclusions

The current study provides preclinical data supporting the notion of sex differences in the mechanisms underlying the development of posttraumatic headache. The data indicates that females develop posttraumatic cephalic pain hypersensitivity for a longer duration, as well as extended latent hypersensitivity to the headache trigger GTN. In addition, our data suggests that the acute PTH-like pain symptoms in females are CGRP-independent, and that the latent sensitization to GTN involves CGRP signaling to a much lesser extent than in males. The observed sex difference in pain response and CGRP involvement may have implication for the mechanisms that link female sex to the increased propensity to develop PTH, and potentially its therapeutic approaches.

